# Interplay between the cell envelope and mobile genetic elements shapes gene flow in populations of a nosocomial pathogen

**DOI:** 10.1101/2020.12.09.417816

**Authors:** Matthieu Haudiquet, Amandine Buffet, Olaya Rendueles, Eduardo P.C. Rocha

## Abstract

Mobile genetic elements (MGEs) drive genetic transfers between bacteria using mechanisms that are affected by the cell envelope composition, notably the capsule. Here, we show that capsules constrain phage-mediated gene flow between closely related serotypes in *Klebsiella pneumoniae*, a high-priority nosocomial enterobacteria. Serotype-specific phage pressure may also explain the inactivation of capsule genes, which occur frequently and recapitulate the capsule biosynthetic pathway. We show that plasmid conjugation is increased upon capsule inactivation and that capsule re-acquisition leaves long recombination tracts around the capsular locus. This suggests that capsule inactivation by phage pressure facilitates its subsequent re-acquisition by conjugation, a process re-wiring gene flow towards novel lineages whenever it leads to serotype swaps. These results reveal the basis of trade-offs between the evolution of virulence and multidrug resistance. They also caution that some alternatives to antibiotic therapy may select for capsule inactivation, thus decreasing virulence but facilitating antibiotic resistance genes acquisition.

## INTRODUCTION

Mobile genetic elements (MGE) drive horizontal gene transfer (HGT) between bacteria, which may result in the acquisition of virulence factors and antibiotic resistance genes (*1, 2*). DNA can be exchanged between cells via virions or conjugative systems (*3, 4*). Virions attach to specific cell receptors to inject their DNA into the cell, which restricts their host range (*5*). When replicating, bacteriophages (henceforth phages) may package bacterial DNA and transfer it across cells (transduction). Additionally, temperate phages may integrate into the bacterial genome as prophages, eventually changing the host phenotype (*4*). In contrast, DNA transfer by conjugation involves mating-pair formation (MPF) between a donor and a recipient cell (*6*). Even if phages and conjugative elements use very different mechanisms of DNA transport, both depend crucially on interactions with the cell envelope of the recipient bacterium. Hence, changes in the bacterial cell envelope may affect their rates of transfer.

*Klebsiella pneumoniae* (Kpn) is a gut commensal that has become a major threat to public health (*7, 8*), and is acquiring MGEs encoding antibiotic resistance (ARG) and virulence factors at a fast pace (*2, 9*). This propensity is much higher in epidemic nosocomial multi-drug resistant lineages than in hypervirulent strains producing infections in the community (*10*). Kpn is a particularly interesting model system to study the interplay between HGT and the cell envelope because it is covered by a nearly ubiquitous Group I (or Wzx/Wzy-dependent) polysaccharide capsular structure (*11, 12*), which is the first point of contact with incoming MGEs. Similar capsule loci are present in many bacteria (*13*). There is one single capsule locus in Kpn (*14*), which evolves quickly by horizontal transfer and recombination (*16, 17*). It contains a few conserved genes encoding the proteins necessary for the multi-step chain of assembly and exportation, which flank a highly variable region encoding enzymes that determine the oligosaccharide combination, linkage and modification (and thus the serotype) (*18*). There are more than 140 genetically distinct capsular locus types (CLT), of which 76 have well characterized chemical structures and are referred to as serotypes (*18*). Kpn capsules can extend well beyond the outer membrane, up to 420nm, which is 140 times the average size of the peptidoglycan layer (*19*). They enhance cellular survival to bacteriocins, immune response, and antibiotics (*20*–*22*), being a major virulence factor of the species. Intriguingly, the multi-drug resistant lineages of Kpn exhibit higher capsular diversity than the virulent ones, which are almost exclusively of the serotype K1 and K2 (*10*).

By its size, the capsule hides phage receptors and can block phage infection (*23*). Since most Kpn are capsulated, many of its virulent phages evolved to overcome the capsule barrier by encoding serotype-specific depolymerases in their tail proteins (*24, 25*). For the same reason, phages have evolved to use the capsule for initial adherence before attaching to the primary cell receptor. Hence, instead of being hampered by the capsule, many Kpn phages have become dependent on it (*26, 27*). This means that the capsule may affect the rates of HGT positively or negatively depending on how it enables or blocks phage infection. Furthermore, intense phage predation may select for capsule swap or inactivation, because this renders bacteria resistant to serotype-specific phages. While such serotype swaps may allow cells to escape phages to which they were previously sensitive, albeit exposing them to new infectious phages, capsule inactivation can confer pan-resistance to capsule-dependent phages (*26*). In contrast, very little is known on the effect of capsules on conjugation, except that it is less efficient between a few different serotypes of *Haemophilus influenzae* (*28*). The interplay between phages and conjugative elements and the capsule has the potential to strongly impact Kpn evolution in terms of both virulence and antibiotic resistance because of the latter’s association with specific serotypes and MGEs.

The capsule needs MGEs to vary by HGT, but may block the acquisition of the very same MGEs. Moreover, capsulated species are associated with higher rates of HGT (*29*). There is thus the need to understand its precise impact on gene flow and how the latter affects capsule evolution. Here, we leverage a very large number of genomes of Kpn to investigate these questions using computational analyses that are complemented with experimental data. As a result, we propose a model of capsule evolution involving loss and re-gain of function. This model explains how the interplay of the capsule with different MGEs can either lower, increase or re-wire gene flow depending on the way capsule affects their mechanisms of transfer.

## Results

### Gene flow is higher within than between serotype groups

We reasoned that if MGEs are specifically adapted to serotypes, then genetic exchanges should be more frequent between bacteria of similar serotypes. We used Kaptive (*18*) to predict the CLT in 3980 genomes of Kpn. Around 92% of the isolates could be classed with good confidence-level. They include 108 of the 140 previously described CLTs of *Klebsiella spp*. The pangenome of the species includes 82,730 gene families, which is 16 times the average genome. It contains 1431 single copy gene families present in more than 99% of the genomes that were used to infer a robust rooted phylogenetic tree of the species (average ultra-fast bootstrap of 98%, Figure 1A). Rarefaction curves suggest that we have extensively sampled the genetic diversity of Kpn genomes, its CLTs, plasmids and prophages (Figure 1B). We then inferred the gains and losses of each gene family of the pangenome using PastML and focused on gene gains in the terminal branches of the species tree predicted to have maintained the same CLT from the node to the tip (91% of branches). This means that we can associate each of these terminal branches to one single serotype. We found significantly more genes acquired (co-gained) in parallel by different isolates having the same CLT than expected by simulations assuming random distribution in the phylogeny (1.95x, Z-test p<0.0001, Figure 1C). This suggests that Kpn exhibits more frequent within-serotype than between-serotype genetic exchanges.

**Figure 1.**
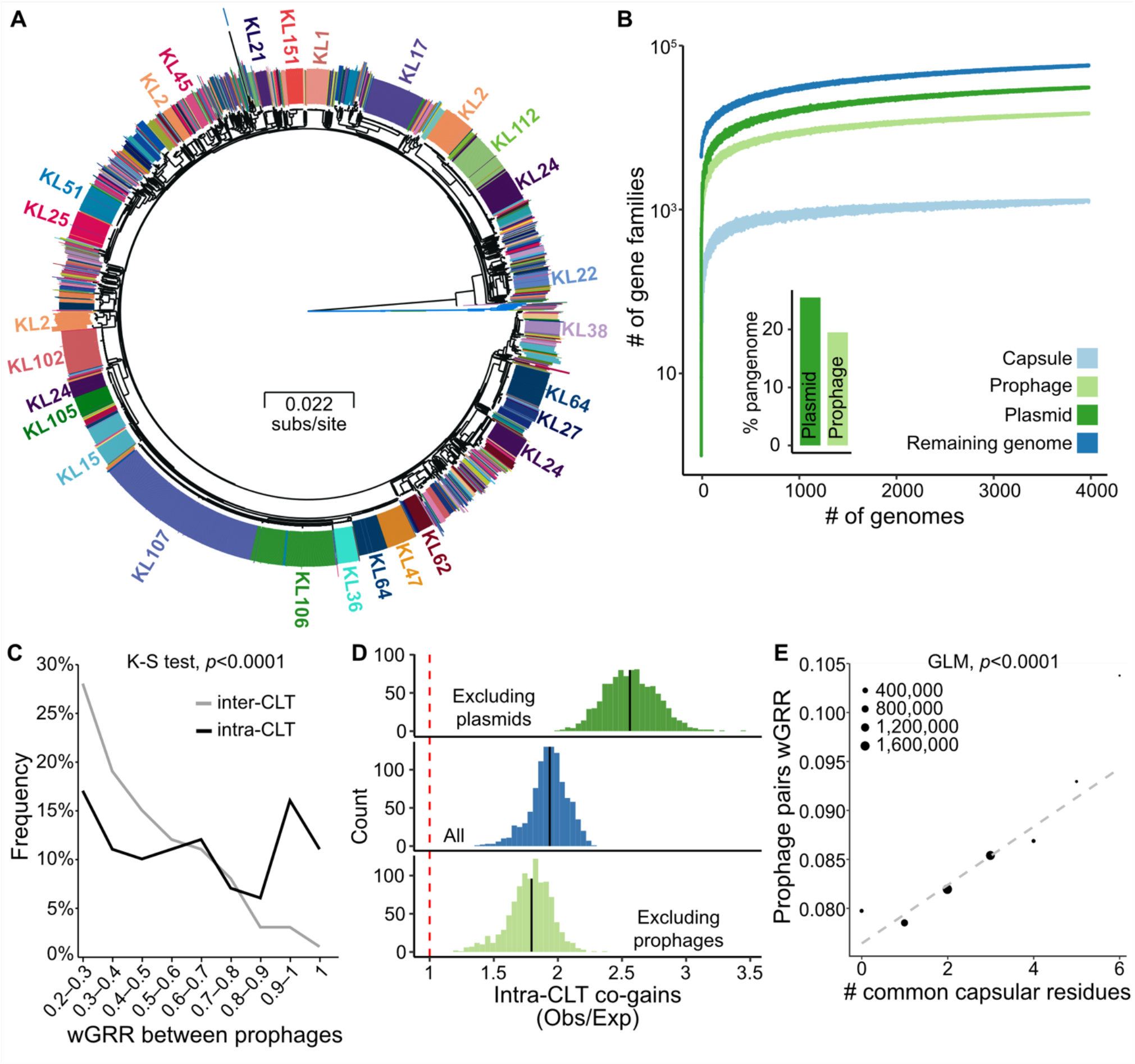
Gene flow is higher between strains of the same serotype. **A**. Phylogenetic tree with the 22 *Klebsiella quasipneumoniae* subsp. *similipneumoniae* (Kqs) strains as an outgroup (blue branches). The annotation circle represents the 108 CLTs predicted by Kaptive. The largest clusters of CLTs (>20 isolates) are annotated (full list in dataset SD1). **B**. Rarefaction curves of the pangenome of prophages, plasmids, capsule genes and all remaining genes (Genome). The points represent 50 random samples for each bin (bins increasing by 10 genomes). The inset bar plot represents the percentage of gene families of the Kpn pangenome including genes of plasmids or prophages. **C**. Gene repertoire relatedness between independently acquired prophages (for wGRR>0.2) in bacteria of different (inter-CLT, grey) or identical CLT (intra-CLT, black). **D**. Histogram of the excess of intra-CLT co-gains in relation to those observed inter-CLT (Observed/expected ratio obtained by 1000 simulations). The analysis includes all genes (center), excludes prophages (bottom), or excludes plasmids (top). **E**. Linear regression of the wGRR between pairs of prophages and the number of capsular residues in common between their hosts. The points represent the mean for each category, with their size corresponding to the number of pairs per category, but the regression was performed on the original data which is composed of several millions of pairs.

Given the tropism of Kpn phages to specific serotypes, we wished to clarify if phages contribute to the excess of intra-CLTs genetic exchanges. Since transduction events cannot be identified unambiguously from the genome sequences, we searched for prophage acquisition events, i.e. for the transfer of temperate phages from one bacterial genome to another. We found that 97% of the strains were lysogens, with 86% being poly-lysogens, in line with our previous results in a much smaller dataset (*26*). In total, 9886 prophages were identified in the genomes, with their 16,319 gene families accounting for 19.5% of the species pan-genome (Figure 1B). We then measured the gene repertoire relatedness weighted by sequence identity (wGRR) between all pairs of prophages. This matrix was clustered, resulting in 2995 prophage families whose history of vertical and horizontal transmissions was inferred using the species phylogenetic tree (see Methods). We found 3269 independent infection events and kept one prophage for each of them. We found that pairs of independently infecting prophages are 1.7 times more similar when in bacteria with identical rather than different CLTs (Figure 1C Two-sample Kolmogorov-Smirnov test, p<0.0001). To confirm that phage-mediated HGT is favoured between strains of the same CLT, we repeated the analysis of gene co-gains after removing the prophages from the pangenome. As expected, the preference toward same-CLT exchanges decreased from 1.95x to 1.73x (Figure 1D). This suggests that HGT tends to occur more frequently between strains of similar serotypes than between strains of different serotypes, a trend that is amplified by the transfer of temperate phages.

Most of the depolymerases that allow phages to overcome the capsule barrier act on specific di- or trisaccharide, independently of the remaining monomers (*30*–*32*). This raises the possibility that phage-mediated gene flow could be higher between strains whose capsules have common oligosaccharide residues. To test this hypothesis, we compiled the information on the 76 capsular chemical structures described (*33*). The genomes with these CLTs, 59% of the total, show a weak but significant proportionality between prophage similarity and the number of similar residues in their host capsules (Figure 1E), i.e. prophages are more similar between bacteria with more biochemically similar capsules.

### Recombination swaps biochemically-related capsules

To understand the genetic differences between serotypes and how these could facilitate swaps, we compared the gene repertoires of the different capsular loci (between *galF* and *ugd*, Figure S1). As expected from previous works (*11, 18*), this analysis revealed a clear discontinuity between intra-CLT comparisons that had mostly homologous genes and the other comparisons, where many genes (average=10) lacked homologs across serotypes (Wilcoxon test, p<0.0001, Figure 2A). As a result, the capsule pangenome contains 325 gene families that are specific to a CLT (out of 547, see Methods). This implicates that serotype swaps require the acquisition of multiple novel genes by horizontal transfer. To quantify and identify these CLT swaps, we inferred the ancestral CLT in the phylogenetic tree and found a rate of 0.282 swaps per branch (see Methods). We then identified 103 highly confident swaps, some of which occurred more than once (Figure 2B). We used the chemical characterization of the capsules described above to test if it could explain these results. Indeed, swaps occurred between capsules with an average of 2.42 common sugars (mean Jaccard similarity 0.54), more than the average value across all other possible CLT pairs (1.98, mean Jaccard similarity 0.38, Wilcoxon test, p<0.0001, Figure S2A). Interestingly, the wGRR of the swapped loci is only 3% higher than the rest of pairwise comparisons (Figure S2B). This suggests that successful swaps are poorly determined by the differences in gene repertoires. Instead, they are more frequent between capsules that have more similar chemical composition.

**Figure 2.**
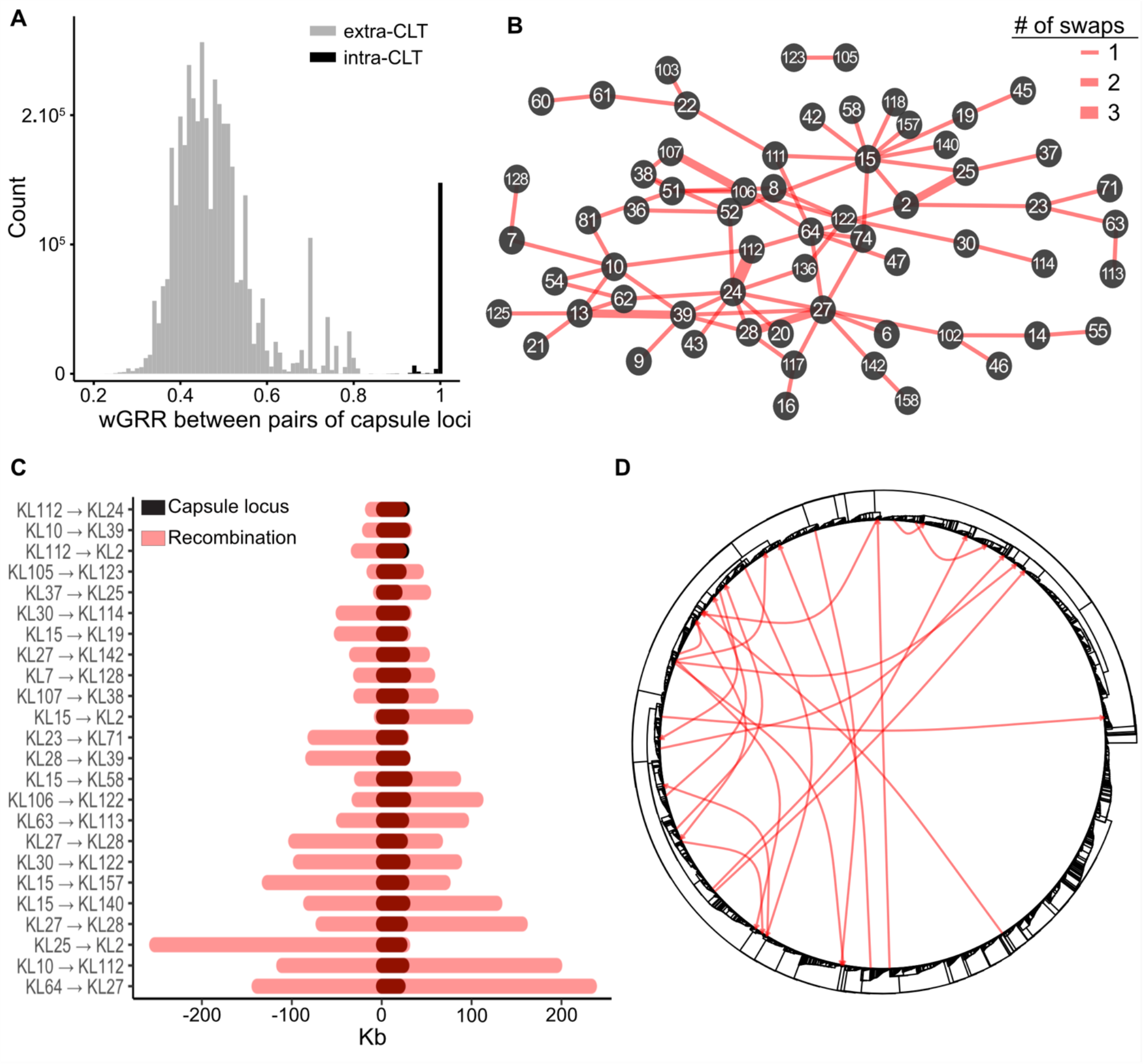
Homologous recombination events lead to frequent CLT swaps. **A**. Histogram of the comparisons of gene repertoire relatedness (wGRR) between capsular loci of the same (intra-) or different (inter-) CLT. **B**. Network of CLT swaps identified by ancestral state reconstruction, with edge thickness corresponding to the number of swaps, and numbers **x** within nodes corresponding to the CLT (KL**x**). **C**. Recombination encompassing the capsule locus detected with Gubbins. The positions of the tracts are represented in the same scale, where the first base of the *galF* gene was set as 0. **D**. Putative donor-recipient pairs involved in the CLT swaps of panel C indicated in the Kpn tree.

The existence of a single capsule locus in Kpn genomes suggests that swaps occur by homologous recombination at flanking conserved sequences (*11*). We used Gubbins (*34*) to detect recombination events in the 25 strains with terminal branch serotype swaps and closely-related completely assembled genomes (see Methods). We found long recombination tracts encompassing the capsule locus in 24 of these 25 genomes, with a median length of 100.3kb (Figure 2C). At least one border of the recombination tract was less than 3kb away from the capsule locus in 11 cases (46%). Using sequence similarity to identify the origin of the transfer, we found that most recombination events occurred between distant strains and no specific clade (Figure 2D). We conclude that serotype swaps occur by recombination at the flanking genes with DNA from genetically distant isolates but chemically related capsules.

### Capsule inactivation follows specific paths, might be driven by phage predation and spurs HGT

We sought to investigate whether the aforementioned swaps occur by an intermediary step where cells have inactivated capsule loci, e.g. resulting from abiotic or biotic selective pressures for capsule loss. To do so, we first established the frequency of inactivated capsular loci. We used the Kaptive software to detect missing genes, expected to be encoded in capsule loci found on a single contig. We also used the Kaptive database of capsular proteins to detect pseudo-genes using protein-DNA alignments in all genomes. We found 55 missing genes and 447 pseudogenes, among 9% of the loci. The frequency of pseudogenes was not correlated with the quality of the genome assembly (see Methods), and all genomes had at least a part of the capsule locus. We classed 11 protein families as essential for capsule production (Table S1). At least one of these essential genes was missing in 3.5% of the loci, which means these strains are likely non-capsulated (Figure 3A). These variants are scattered in the phylogenetic tree with no particular clade accounting for the majority of these variants, e.g. there are non-capsulated strains in 61 of the 617 sequence types (ST) identified by Kleborate. These results suggest that capsule inactivation has little phylogenetic inertia, i.e. it’s a trait that changes very quickly, either because the variants are counter-selected or because capsules are quickly re-acquired. Accordingly, the Pagel’s Lambda test (*35*) showed non-significant phylogenetic inertia (p>0.05). Hence, capsule inactivation is frequent but non-capsulated lineages do not persist for long periods of time.

**Figure 3.**
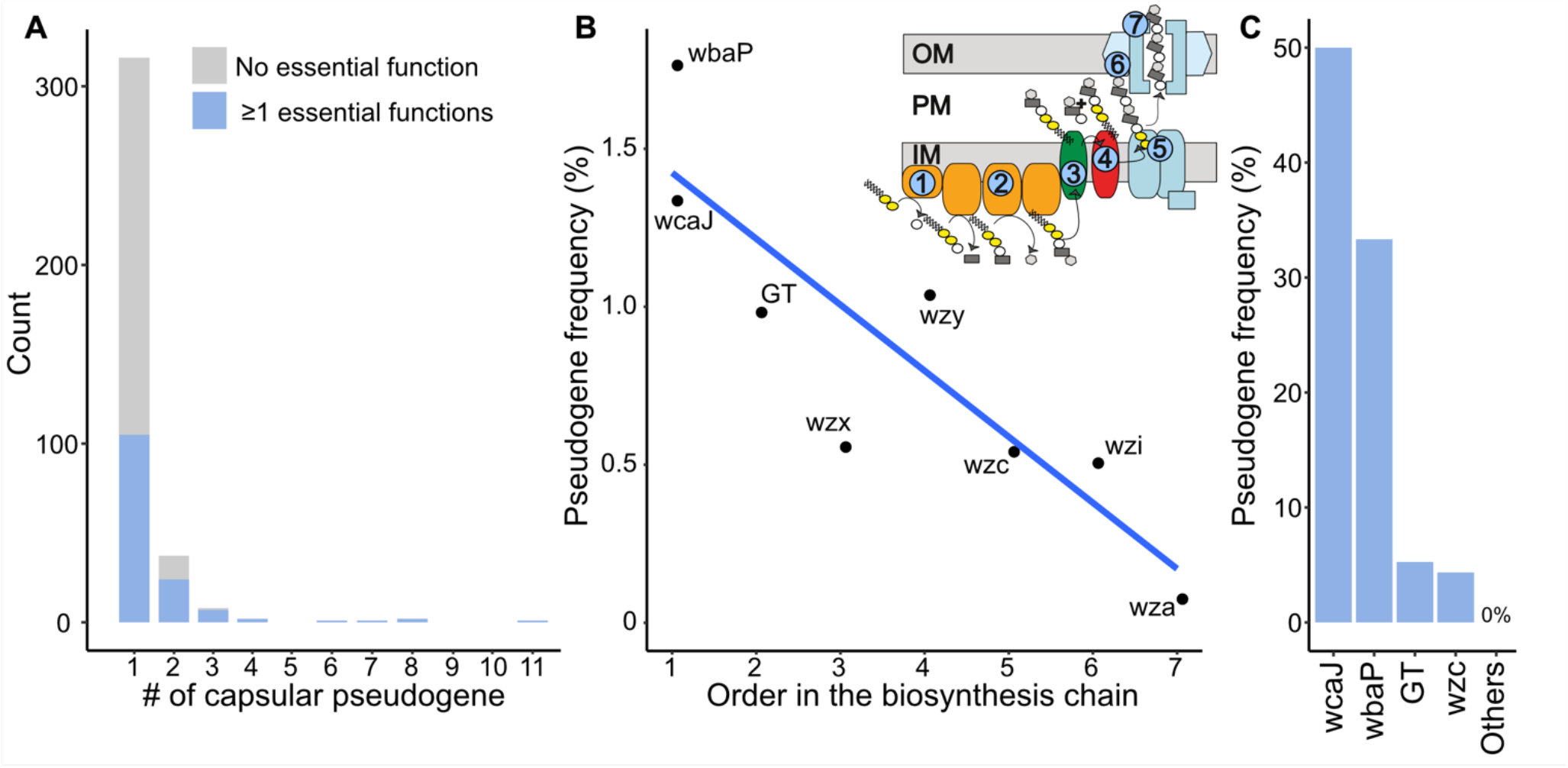
Loss of function in the capsule locus. **A**. Distribution of the number of capsular pseudogenes per genome, split in two categories: loci lacking a functional essential capsular gene (blue, non-capsulated strains), and other loci lacking non-essential capsular genes (white, not categorized as non-capsulated strains). **B**. Linear regression between the pseudogene frequency and the rank of each gene in the biosynthesis pathway (p=0.005, R^2^=0.75). The numbers in the scheme of the capsule assembly correspond to the order in the biosynthesis pathway. **C**. Frequency of pseudogenes arising in the non-capsulated clones isolated in eight different strains after *ca*. 20 generations in LB growth medium. Genomes containing several missing genes and pseudogenes are not included.

We further investigated the genetic pathways leading to capsule inactivation. Interestingly, we found that the pseudogenization frequency follows the order of biosynthesis of the capsule (Linear regression, p=0.005, R^2^=0.75), with the first (*wbaP* or *wcaJ*) and second step (Glycosyl-transferases, GT) being the most commonly inactivated when a single essential gene is a pseudogene (Figure 3B). The overall frequency of gene inactivation drops by 14% per rank in the biosynthesis chain. To test if similar results are found when capsules are counter-selected in the laboratory, we analysed a subset of populations stemming from a short evolution experiment in which different strains of *Klebsiella spp*. were diluted daily during three days (*ca*. 20 generations) under agitation in LB, a medium known to select for capsule inactivation (*36*). After three days, non-capsulated clones emerged in 22 out of 24 populations from eight different ancestral strains. We isolated one non-capsulated clone for Illumina sequencing from each population and searched for the inactivating mutations. We found that most of these were localized in *wcaJ* and *wbaP* (Figure 3C). In accordance with our comparative genomics analysis, we found fewer loss-of-function mutations in GTs and *wzc* and none in the latter steps of the biosynthetic pathway. These results strongly suggest that mutations leading to the loss of capsule production impose a fitness cost determined by the position of the inactivated gene in the biosynthesis pathway.

Once a capsule locus is inactivated, the function can be re-acquired by: 1) reversion mutations fixing the broken allele, 2) restoration of the inactivated function by acquisition of a gene from another bacterium, eventually leading to a chimeric locus, 3) replacement of the entire locus leading to a CLT swap. Our analyses of pseudogenes provide some clues on the relevance of the three scenarios. We found 111 events involving non-sense point mutations. These could eventually be reversible (scenario 1) if the reversible mutation arises before other inactivating changes accumulate. We also observed 269 deletions (100 of more than 2 nucleotides) in the inactivated loci. These changes are usually irreversible in the absence of HGT. We then searched for chimeric loci (scenario 2), i.e. CLTs containing at least one gene from another CLT. We found 35 such loci, accounting for *ca*. 0.9% of the dataset (for example a *wzc_KL1* allele in an otherwise KL2 loci), with only one occurrence of a *wcaJ* allele belonging to another CLT, and none for *wbaP*. Finally, the analysis of recombination tracts detailed above reveals frequent replacement of the entire locus between *galF* and *ugd* by recombination (Figure S1, scenario 3).

Since re-acquisition of the capsule function might often require HGT, we enquired if capsule inactivation was associated with higher rates of HGT. Indeed, the number of genes gained by HGT per branch of the phylogenetic tree is higher in branches where the capsule was inactivated than in the others (Two-sample Wilcoxon test, p<0.0001, Figure 4A), even if these branches have similar sizes (Figure S4). We compared the number of phages and conjugative systems acquired in the branches where capsules were inactivated against the other branches. This revealed significantly more frequent (3.6 times more) acquisition of conjugative systems (Fisher’s exact test, p<0.0001, Figure 4B) upon capsule inactivation. This was also the case, to a lesser extent, for prophages. Intriguingly, we observed even higher relative rates of acquisition of these MGEs in branches where the serotype was swapped (Figure 4A,B), but in this case, the acquisition of prophages was more frequent than that of conjugative systems (6.5 vs. 4.5 times more). However, branches where capsules were swapped are 2.7 times longer than the others, precluding strong conclusions (Figure S4). Overall, periods of capsule inactivation are associated with an excess of HGT. This facilitates the re-acquisition of a capsule and the novel acquisition of other potentially adaptive traits.

**Figure 4.**
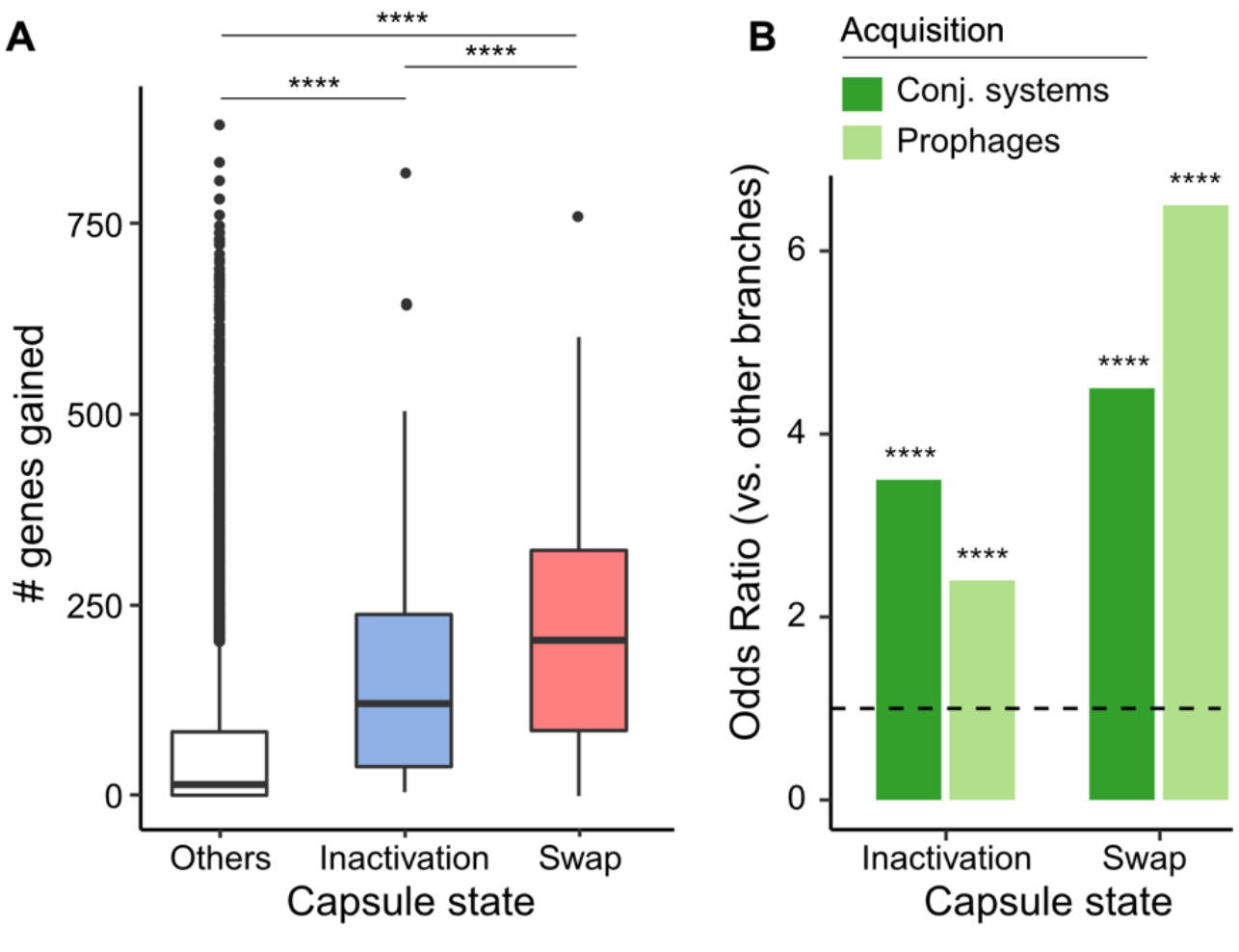
Changes in capsule state impact HGT. **A**. Number of genes gained in branches of the phylogenetic tree where the capsule was inactivated, swapped and in the others (Two-samples Wilcoxon test). **B**. Increase in the frequency of acquisition of prophages and conjugative systems on branches where the capsule was either inactivated or swapped, relative to the other branches, as represented by odds-ratio (Fisher’s exact test). ****: p-value<0.0001

### Conjugative systems are frequently transferred across serotypes

The large size of the Kpn capsule locus is difficult to accommodate in the phage genome and the tendency of phages to be serotype-specific makes them unlikely vectors of novel capsular loci. Also, the recombination tracts observed in Figure 2C are too large to be transduced by most temperate phages of Kpn, whose prophages average 46 kb (*26*). Since Kpn is not naturally transformable, we hypothesized that conjugation is the major driver of capsule acquisition. Around 80% of the strains encode a conjugative system and 94.4% have at least one plasmid, the latter alone making 25.5% of the pan-genome (Figure 1B). We estimate that 41% of the conjugative systems in Kpn are not in plasmids but in integrative conjugative elements (ICE). Since ICEs and conjugative plasmids have approximately similar sizes (*37*), the joint contribution of ICEs and plasmids in the species pangenome is very large.

We identified independent events of infection by conjugative systems as we did for prophages (see above). The 5144 conjugative systems fell into 252 families with 1547 infection events. On average, pairs of conjugative systems acquired within the same CLT were only 3% more similar than those in different CLTs. This suggests an opposite behaviour of phage- and conjugation-driven HGT, since the former tend to be serotype-specific whereas the latter are very frequently transferred across serotypes. This opposition is consistent with the analysis of co-gains (Figure 1D), which were much more serotype-dependent when plasmids were excluded from the analysis and less serotype-dependent when prophages were excluded. To further test our hypothesis, we calculated the number of CLTs where one could find each family of conjugative systems or prophages and then compared these numbers with the expectation if they were distributed randomly across the species. The results show that phage families are present in much fewer CLTs than expected, whereas there is no bias for conjugative systems (Figure 5). We conclude that conjugation spreads plasmids across the species regardless of serotype. Together, these results reinforce the hypothesis that conjugation drives genetic exchanges between strains of different serotypes, decreasing the overall bias towards same-serotype exchanges driven by phages.

**Figure 5.**
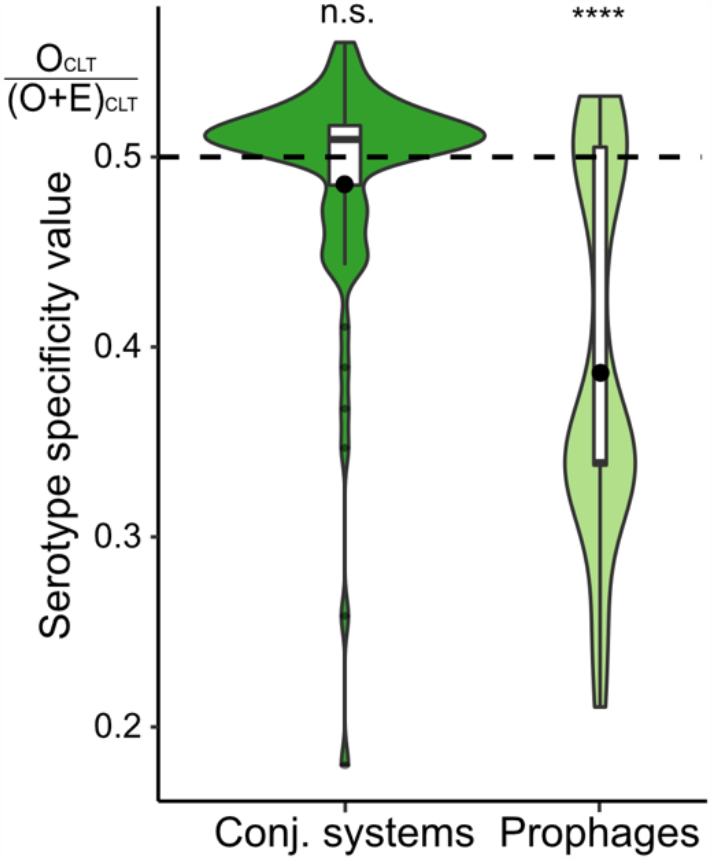
Serotype specificity of prophages and conjugative systems. O_CLT_: Observed number of serotypes infected per family of homologous element. E_CLT_: expected number of serotypes infected per family of homologous element generated by 1000 simulations (see Methods). When the elements distribute randomly across serotypes, the value is 0.5 (dashed line). Very low values indicate high serotype-specificity. One-sample Wilcoxon test. ****: p-value<0.0001

### Capsule inactivation results in increased conjugation efficiency

Together, these elements led us to hypothesize that capsule inactivation results in higher rates of conjugation. This is consistent with the observation that terminal branches associated with inactive capsules have a higher influx of conjugative systems than prophages (Figure 4B). In the absence of published data on the frequency of conjugation in function of the presence of a capsule, we tested experimentally our hypothesis on a diverse set of *Klebsiella* isolates composed of four strains from different STs: three *Klebsiella pneumoniae sensu stricto* (serotypes K1 and K2) and one *Klebsiella variicola* (serotype K30, also found in Kpn). To test the role of the capsule in plasmid acquisition, we analysed the conjugation efficiency of these strains and their non-capsulated counterparts, deprived of *wcaJ*, the most frequently pseudogene in the locus both in the genome data and in our experimental evolution (see Methods). For this, we built a plasmid that is mobilized in *trans*, i.e. once acquired by the new host strain it cannot further conjugate. This allows to measure precisely the efficiency of conjugation between the donor and the recipient strain. In agreement with the results of the computational analysis, we found that the efficiency of conjugation is systematically and significantly higher in the mutant than in the associated WT for all four strains (paired Wilcoxon test, p-value=0.002, Figure 6). On average, non-capsulated strains conjugated 8.06 times more than capsulated strains. Interestingly, the magnitude of the difference in conjugation rates is inversely proportional to the wild type frequency, possibly because some strains already conjugate at very high rates even with a capsule. These experiments show that the ability to receive a conjugative element is increased in the absence of a functional capsule. Hence, non-capsulated variants acquire more genes by conjugation than the others. Interestingly, if the capsule is transferred by conjugation, this implies that capsule inactivation favours the very mechanism leading to its subsequent re-acquisition.

**Figure 6.**
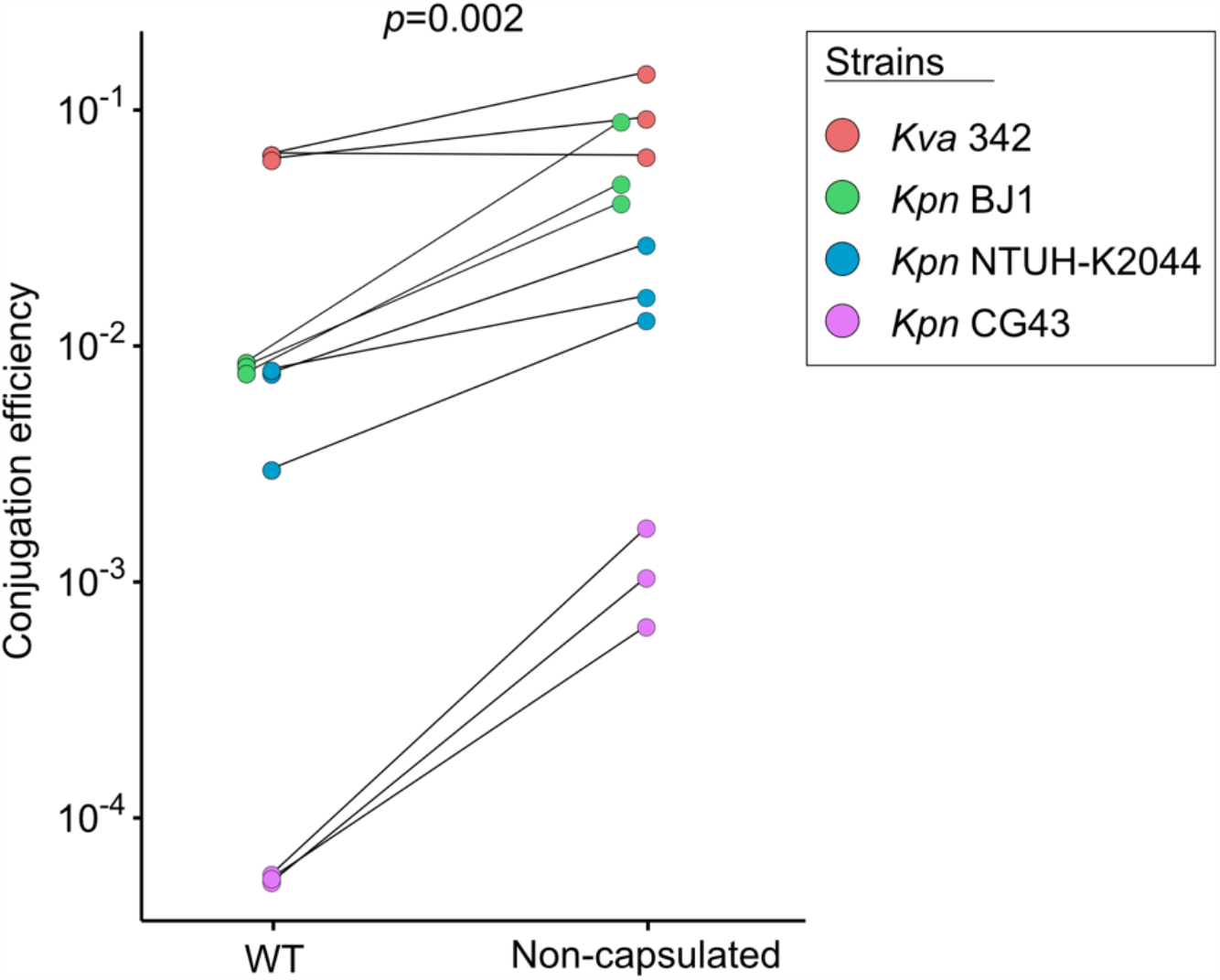
Capsules negatively impact conjugation. Conjugation efficiency of wild type (WT) and their associated non-capsulated (*ΔwcaJ*) mutants. The conjugation efficiency is represented on a log-scale. Each pair of points represents a biological replicate. The p-value for the paired Wilcoxon test is displayed.

## Discussion

The specificity of many Kpn phages to one or a few chemically related serotypes is presumably caused by their reliance on capsules to adsorb to the cell surface and results from the longstanding co-evolution of phages with their Kpn hosts. One might invoke environmental effects to explain these results, since populations with closely related serotypes might often co-occur and thus potentiate more frequent cross-infections. However, the same ecological bias would be expected for conjugation and this could not be detected. Instead, phages infecting bacteria with related serotypes might carry capsule depolymerases, which are known to act on di- or tri-saccharides (*30*–*32*) similar across these serotypes. This fits our previous studies on the infection networks of Kpn prophages (*26*) and suggests that a population of cells encoding and expressing a given serotype has more frequent phage-mediated genetic exchanges with bacteria of identical or similar serotypes (Figure 6A). In this context, phages carrying multiple capsule depolymerases have broader host range and may have a key role in phage-mediated gene flow between very distinct serotypes. For example, one broad host range virulent phage has been found to infect ten distinct serotypes because it encodes an array of at least nine depolymerases (*25*).

The interplay between the capsule and conjugative elements has been much less studied. Our comparative genomics analyses reveal that conjugation occurs across the species independently of the capsule serotypes. Furthermore, our experimental data shows that non-capsulated bacteria are up to 20 times more receptive to plasmid conjugation than the other bacteria, an effect that seems more important for wild-type capsulated bacteria that are poor recipients. These results are likely to be relevant not only for non-capsulated strains, but also for those not expressing the capsule at a given moment. If so, repression of the expression of the capsule may allow bacteria to escape phages and endure extensive acquisition of conjugative elements. These results may also explain a longstanding conundrum in Kpn. The hypervirulent lineages of Kpn, which are almost exclusively of serotypes K1 and K2 (*10, 38*), have reduced pangenome, plasmid and capsule diversity. They also often carry additional factors like *rmpA* upregulating the expression of the capsule (*38*). In contrast, they are very rarely multi-drug resistant. Our data suggests that the protection provided by thick capsules hampers the acquisition of conjugative elements, which are the most frequent vectors of antibiotic resistance. Furthermore, the moments of capsule swap or inactivation are expected to be particularly deleterious for hyper-virulent clones, because the capsule is a virulence factor, thus further hampering their ability to acquire the conjugative MGEs that carry antibiotic resistance. This may have favoured a specialization of the clones into either hyper-virulent or multi-drug resistance. Unfortunately, the capsule is not an insurmountable barrier for conjugation, and recent reports have uncovered the emergence of worrisome multi-drug resistant hyper-virulent clones (*39, 40*).

We observed that branches of isolates lacking a functional capsule have higher rates of acquisition of conjugative systems than prophages, whereas those where there was a capsule swap have the inverse pattern (Figure 4B). What could justify these differences in the interplay of the capsule with phages and conjugative elements? Phages must adsorb on the cell surface, whereas there are no critical positive determinants for incoming conjugation pilus (*41*). As a result, serotypes swaps may affect much more the flow of phages than that of conjugative elements. Capsule loss may have an opposite effect on phages and plasmids: it removes a point of cell attachment for phages, decreasing their infection rates, and removes a barrier to the conjugative pilus, increasing their ability to transfer DNA. Hence, when a bacterium loses a capsule, e.g. because of phage predation, it becomes more permissive for conjugation. In contrast, when a bacterium acquires a novel serotype it may become sensitive to novel phages resulting in rapid turnover of its prophage repertoire. These results implicate that conjugation should be much more efficient at spreading traits across the entire Kpn species than phage-mediated mechanisms, which could have an important role for intra-serotype HGT (Figure 7A).

**Figure 7.**
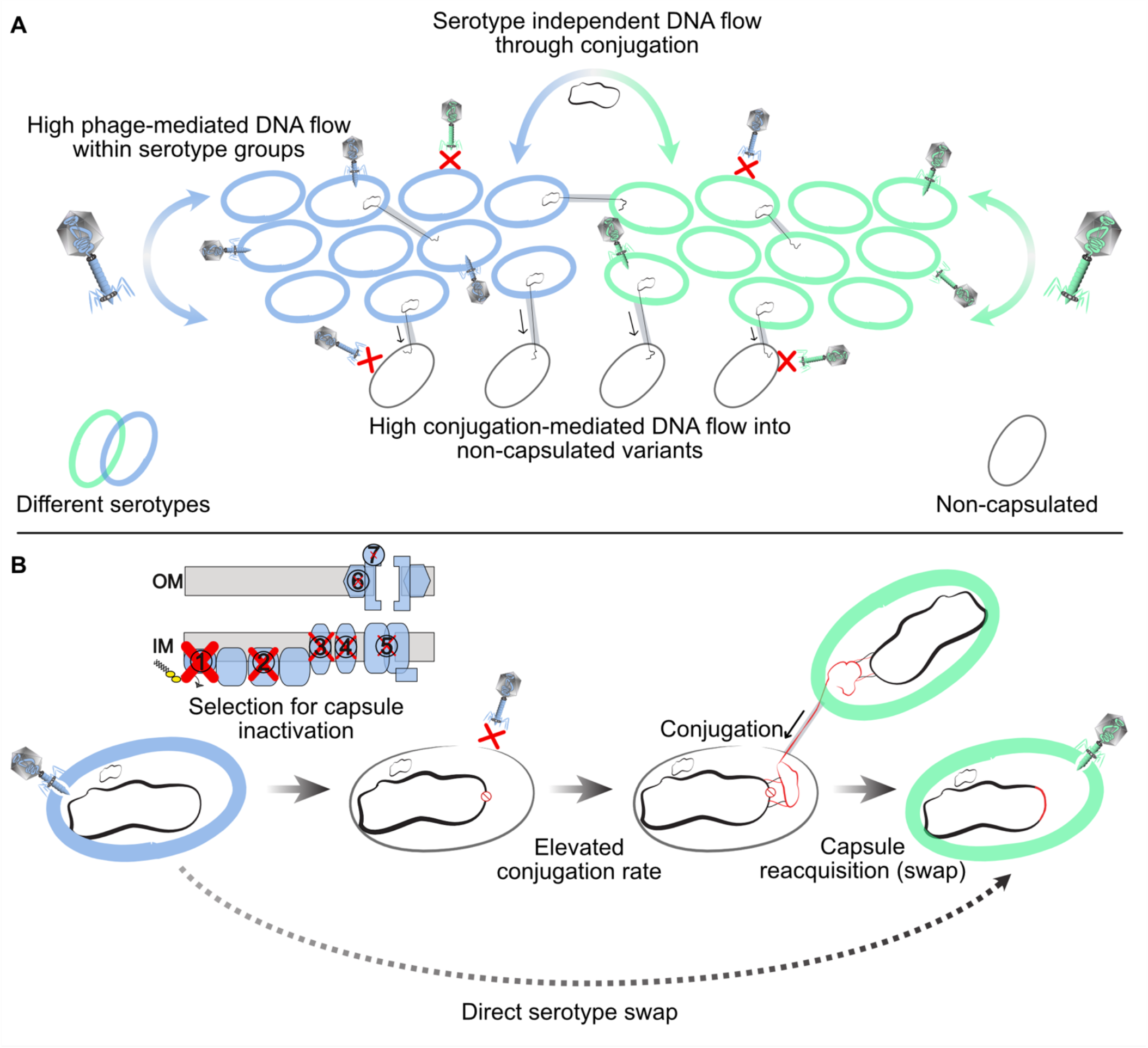
A model for the interplay between serotypes and mobile elements. **A**. The capsule impacts Kpn gene flow. A bacterial population expressing a given serotype (blue or green) preferentially exchanges DNA by phage-mediated processes with bacteria of identical or similar serotypes. Such flow may be rare towards non-capsulated bacteria because they are often resistant to Kpn phages. In contrast, conjugation occurs across serotypes and is more frequent to non-capsulated bacteria. **B**. A model for serotype swaps in Kpn. Capsule inactivation is occasionally adaptive. The pseudogenization process usually starts by the inactivation of the genes involved in the early stages of the capsule biosynthesis, as represented by the size of the red cross on the capsule assembly scheme. Non-capsulated strains are often protected from Kpn phage infections whilst acquiring more genes by conjugation. This increases the likelihood of capsule re-acquisition. Such re-acquisition can bring a new serotype, often one that is chemically similar to the previous one, and might be driven by conjugation because of its high frequency in non-capsulated strains. Serotype swaps re-wire phage-mediated transfers.

The existence of serotype swaps has been extensively described in the literature for Kpn (*17*) and many other species (*42, 43*). Whether these swaps implicate a direct serotype replacement, or an intermediate non-capsulated state, is not sufficiently known. Several processes are known to select for capsule loss in some bacteria, including growth in rich medium (*36*), phage-pressure, and immune response (*26, 27, 44, 45*) (Figure 7B). Because of the physiological effects of these losses, and their impact on the rates and types of HGT, it’s important to quantify the frequency of inactivated (or silent) capsular loci and the mechanisms favouring it. Our study of pseudogenization of capsular genes revealed a few percent of non-capsulated strains scattered in the species tree, opening the possibility that non-capsulated strains are a frequent intermediate step of serotype swap. The process of capsule inactivation is shaped by the capsule biosynthesis pathway, the frequency of pseudogenization decreasing linearly with the rank of the gene in the capsule biosynthesis pathway. This suggests a major role for epistasis in the evolutionary pathway leading to non-capsulated strains. Notably, the early inactivation of later genes in the biosynthesis pathway, while the initial steps are still functional, can lead to the sequestration of key molecules at the cell envelope or the toxic accumulation of capsule intermediates (Figure 7B). Accordingly, Δ*wza* and Δ*wzy* mutants, but not Δ*wcaJ*, lead to defects in the cell envelope of the strain Kpn SGH10 (*46*). Capsule re-acquisition is more likely driven by conjugation than by phages. Hence, the increased rate of acquisition of conjugative elements by non-capsulated strains may favour the process of capsule re-acquisition. Cycles of gain and loss of capsular loci have been previously hypothesized in the naturally transformable species *Streptococcus pneumoniae*, where vaccination leads to counterselection of capsulated strains and natural transformation seems to increase recombination in non-capsulated clades (*47*). Tracts encompassing the capsule locus were found in serotype-swapped *S. pneumoniae* isolates (*48*), although they were smaller to those found in Kpn (median length 42.7kb).

Our results are relevant to understand the interplay between the capsule and other mobile elements in Kpn or other bacteria. We expect to observe more efficient conjugation when the recipient bacteria lack a capsule in other species. For example, higher conjugation rates of non-capsulated strains may help explain their higher recombination rates in *Streptococcus* (*49*). The serotype-specificity of phages also opens intriguing possibilities for them and for virion-derived elements. For example, some *Escherichia coli* strains able to thrive in freshwater reservoirs have capsular loci acquired in a single-block horizontal transfer from Kpn (*50*). This could facilitate inter-species phage infections (and phage-mediated HGT), since these strains may now be adsorbed by Kpn phages. Gene transfer agents (GTA) are co-options of virions for intra-species HGT that are frequent among alpha-Proteobacteria (*51*) (but not yet described in Kpn). They are likely to have equivalent similar serotype specificity, since they attach to the cell envelope using structures derived from phage tails. Indeed, the infection by the *Rhodobacter capsulatus* GTA model system depends on the host Wzy-capsule (*52*), and non-capsulated variants of this species are phage resistant (*53*) and impaired in GTA-mediated transfer (*54*). Our general prediction is that species where cells tend to be capsulated are going to co-evolve with phages, or phage-derived tail structures, such that the latter will tend to become serotype-specific.

These predictions have an impact in the evolution of virulence and antimicrobial therapy. Some alternatives to antibiotics - phage therapy, depolymerases associated with antibiotics, pyocins, capsular polysaccharide vaccines - may select for the inactivation of the capsule (*26, 44, 55*). Such non-capsulated variants have often been associated with better disease outcomes (*56*), lower antibiotics tolerance (*22*) and reduced virulence (*21*). However, they can also be more successful colonizers of the urinary tract (*57*). Our results suggest that these non-capsulated variants are at higher risk of acquiring resistance and virulence factors through conjugation, because antibiotic resistance genes and virulence factors are often found in conjugative elements in Kpn and in other nosocomial pathogens. Conjugation may also eventually lead to the re-acquisition of functional capsules. At the end of the inactivation-reacquisition process, recapitulated on figure 7, the strains may be capsulated, more virulent and more antibiotic resistant.

## MATERIALS AND METHODS

### Genomes

We used the PanACoTa tool to generate the dataset of genomes (*58*). We downloaded all the 5805 genome assemblies labelled as *Klebsiella pneumoniae sensu stricto* (*Kpn*) from NCBI RefSeq (accessed on October 10^th^ 2018). We removed lower quality assemblies by L90>100. The pairwise genetic distances between all remaining genomes of the species was calculated by order of assembly quality (L90) using MASH (*59*). Strains that were too divergent (MASH distance > 6%) to the reference strain or too similar (<0.0001) to other strains were removed from further analysis. The latter tend to have similar capsule serotypes, and their exclusion does not eliminate serotype swap events. This resulted in a dataset of 3980 strains which were re-annotated with *prokka* (*v1.13.3*) (*60*) to use consistent annotations in all genomes. Erroneous species annotations in the GenBank files were corrected using Kleborate (https://github.com/katholt/Kleborate). This step identified 22 *Klebsiella quasipneumoniae* subspecies *similipneumoniae* (Kqs) genomes that were used to root the species tree and excluded from further analyses. The accession number for each analysed genome is presented in supplementary dataset SD1, along with all the annotations identified in this study.

### Pan- and persistent genome

The pangenome is the full repertoire of homologous gene families in a species. We inferred the pangenome with the connected-component clustering algorithm of MMSeqs2 (*release 5*) (*61*) with pairwise bidirectional coverage > 0.8 and sequence identity > 0.8. The persistent genome was built from the pangenome, with a persistence threshold of 99%, meaning that a gene family must be present in single copy in at least 3940 genomes to be considered persistent. Among the 82,730 gene families of the Kpn pangenome, there were 1431 gene families present in 99% of the genomes, including the Kqs. We used mlplasmids to identify the “plasmid” contigs (default parameters, species “*Klebsiella pneumoniae*”, (*62*)). To identify the pangenome of capsular loci present in the Kaptive database, we used the same method as above, but we lowered the sequence identity threshold to >0.4 to put together more remote homologs.

### Phylogenetic tree

To compute the species phylogenetic tree, we aligned each of the 1431 protein families of the persistent genome individually with mafft (v7.407) (*63*) using the option *FFT-NS-2*, back-translated the sequences to DNA (i.e. replaced the amino acids by their original codons) and concatenated the resulting alignments. We then made the phylogenetic inference using *IQ-TREE* (*v1.6.7.2*) (*64*) using ModelFinder (-m TEST) (*65*) and assessed the robustness of the phylogenetic inference by calculating 1000 ultra-fast bootstraps (-bb 1000) (*66*). There were 220,912 parsimony-informative sites over a total alignment of 1,455,396 bp and the best-fit model without gamma correction was a general time-reversible model with empirical base frequencies allowing for invariable sites (GTR+F+I). We did not use the gamma correction because of branch length scaling issues, which were ten times longer than with simpler models, and is related to an optimization problem with big datasets in IQ-TREE. The tree is very well supported, since the average ultra-fast bootstrap support value was 97.6% and the median 100%. We placed the Kpn species root according to the outgroup formed by the 22 *Kqs* strains. The tree, along with Kleborate annotations, can be visualized and manipulated in https://microreact.org/project/kk6mmVEDfa1o3pGQSCobdH/9f09a4c3

### Capsule locus typing

We used Kaptive (*18*) with default options and the “K locus primary reference” to identify the CLT of strains. We assigned the CLT to “unknown” when the confidence level of Kaptive was “none” or “low” as suggested by the authors of the software. This only represented 7.9% of the genomes.

### Identification of capsule pseudogenes and inactive capsule loci

We first compiled the list of missing expected genes from Kaptive, which is only computed by Kaptive for capsule loci encoded in a single contig. Then we used the Kaptive reference database of Kpn capsule loci to retrieve capsule reference genes for all the identified serotypes. We searched for sequence similarity between the proteins of the reference dataset and the 3980 genome assemblies using blastp and tblastn (v.2.9.0) (*67*). We then searched for the following indications of pseudogenization: stop codons resulting in protein truncation, frameshift mutations, insertions and deletions (supplementary dataset SD2). Truncated and frameshifted coding sequences covering at least 80% of the original protein in the same reading frame were considered functional. Additionally, a pseudogene did not result in a classification of inactivated function if we could identify an intact homolog or analog. For example, if *wcaJ_KL1* has a frameshift, but *wcaJ_KL2* was found in the genome, the pseudogene was flagged and not used to define non-capsulated mutants. Complete gene deletions were identified by Kaptive among capsular loci encoded on a single contig. We built a dictionary of genes that are essential for capsule production by gathering a list of genes (annotated as the gene name in the Kaptive database) present across all CLTs and which are essential for capsule production according to experimental evidence (Table S1). The absence of a functional copy of these essential genes resulted in the classification “non-capsulated” (except *wcaJ* and *wbaP* which are mutually exclusive). To correlate the pseudogenization frequency with the order in the capsular biosynthesis process, we first sought to identify all glycosyl transferases from the different CLTs and grouped them in one category. To do so, we retrieved the GO-molecular functions listed on UniProtKB of the genes within the Kaptive reference database. For the genes that could be ordered in the biosynthesis chain (table S1), we computed their frequency of inactivation by dividing the count of inactivated genes by the total number of times it is present in the dataset.

To test that sequencing errors and contig breaks were not leading to an excess of pseudogenes in certain genomes, we correlated the number of pseudogenes (up to 11) and missing genes with two indexes of sequence quality, namely, the sequence length of the shortest contig at 50% of the total genome length (N50) and the smallest number of contigs whose length sum makes up 90% of genome size (L90). We found no significant correlation in both cases (Spearmans’ correlations, p-values >0.05), suggesting that our results are not strongly affected by sequencing artefacts and assembly fragmentation.

### Genetic similarity

We searched for sequence similarity between all proteins of all prophages or conjugative systems using MMSeqs2 with the sensitivity parameter set to 7.5. The hits were filtered (e-value < 10^−5^, ≥35% identity, coverage > 50% of the proteins) and used to compute the set of bi-directional best hits (BBH) between each genome pair. BBH were used to compute the gene repertoire relatedness between pairs of genomes (weighted by sequence identity):

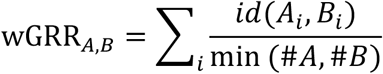

as previously described (*68*), where *A*_*i*_ and *B*_*i*_ is the pair *i* of homologous proteins present in A and B (containing respectively #*A* and #*B* proteins), *id*(*A*_*i*_, *B*_*i*_) is their sequence identity, and min (#*A*, #*B*) is the number of proteins of the element encoding the fewest proteins (#*A* or #*B*). wGRR varies between zero and one. It amounts to zero if there are no BBH between the genomes, and one if all genes of the smaller genome have an identical BBH in the other genome. Hence, the wGRR accounts for both frequency of homology and degree of similarity among homologs.

### Inference of genes ancestral states

We inferred the ancestral state of each pangenome family with PastML (v1.9.23) (*69*) using the maximum-likelihood algorithm MPPA and the F81 model. We also tried to run Count (*70*) with the ML method to infer gene gains and losses from the pangenome, but this took a prohibitive amount of computing time. To check that PastML was producing reliable results, we split our species tree (*cuttree* function in R, package stats) in 50 groups and for the groups that took less than a month of computing time with Count (2500 genomes), we compared the results of Count to those of PastML. The two methods were highly correlated in term of number of inferred gains per branch (Spearman’s correlation test, Rho=0.88, p-value<0.0001). We used the results of PastML, since it was much faster and could handle the whole tree in a single analysis. Since the MPPA algorithm can keep several ancestral states per node if they have similar and high probabilities, we only counted gene gains when both ancestral and descendant nodes had one single distinct state (absent → present).

### Analysis of conjugative systems

To detect conjugative systems, Type IV secretion systems, relaxases, and infer their mating-pair formation (MPF) types, we used TXSScan with default options (*71*). We then extracted the protein sequence of the conjugation systems and used these sequences to build clusters of systems by sequence similarity. We computed the wGRR (see Genetic similarity) between all pairs of systems and clustered them in wGRR families by transitivity when the wGRR was higher than 0.99. This means that some members of the same family can have a wGRR<0.99. This threshold was defined based on the analysis of the shape of the distribution of the wGRR (Figure S3A). We used a reconstruction of the presence of members of each gene family in the species phylogenetic tree to infer the history of acquisition of conjugative elements (see Inference of gene ancestral state). To account for the presence of orthologous families, i.e. those coming from the same acquisition event, we kept only one member of a wGRR family per acquisition event. For example, if a conjugative system of the same family is present in four strains, but there were two acquisition events, we randomly picked one representative system for each acquisition event (in this case, two elements, one per event). Elements that resulted from the same ancestral acquisition event are referred as orthologous systems. We combined the predictions of mlplasmids and TXSScan to separate conjugative plasmids from integrative and conjugative elements (ICEs). The distribution of conjugative system’s mating pair formation (MPF) type in the chromosomes and plasmids is shown in Figure S4.

### Prophage detection

We used PHASTER (*72*) to identify prophages in the genomes (accessed in December 2018). The category of the prophage is given by a confidence score which corresponds to “intact”, “questionable”, or “incomplete”. We kept only the “intact” prophages because other categories often lack essential phage functions. We further removed prophage sequences containing more than three transposases after annotation with ISFinder (*73*) because we noticed that some loci predicted by PHASTER were composed of arrays of insertion sequences. We built clusters of nearly identical prophages with the same method used for conjugative systems. The wGRR threshold for clustering was also defined using the shape of the distribution (Figure S3B). The definition of orthologous prophages follows the same principle than that of conjugative systems, they are elements that are predicted to result from one single past event of infection.

### Serotype swaps identification

We inferred the ancestral state of the capsular CLT with PastML using the maximum-likelihood algorithm MPPA, with the recommended F81 model (*69*). In the reconstruction procedure, the low confidence CLTs were treated as missing data. This analysis revealed that serotype swaps happen at a rate of 0.282 swaps per branch, which are, on average, 0.000218 substitutions/site long in our tree. CLT swaps were defined as the branches where the descendant node state was not present in the ancestral node state. In 92% of the swaps identified by MPPA, there was only one state predicted for both ancestor and descendant node, and we could thus precisely identify the CLT swaps. These swaps were used to generate the network in figure 3A.

### Detection of recombination tracts

We detected recombination tracts with Gubbins v2.4.0 (*34*). Our dataset is too large to build one meaningful whole-genome alignment (WGA). Gubbins is designed to work with closely related strains, so we split the dataset into smaller groups defined by a single ST. We then aligned the genomes of each ST with Snippy v4.3.8 (*74*), as recommended by the authors in the documentation. The reference genome was picked randomly among the complete assemblies of each ST. We analysed the 25 groups in which a CLT swap happened (see above), and for which a complete genome was available as a reference. We launched Gubbins independently for each WGA, using default parameters. We focused on the terminal branches to identify the recombination tracts resulting in CLT swap. We enquired on the origin of the recombined DNA using a sequence similarity approach. We used blastn (*67*) (-task megablast) to find the closest match of each recombination tract by querying the full tract against our dataset of genome assemblies, and mapped the closest match based on the bitscore onto the species tree.

### Identification of co-gains

We used the ancestral state reconstruction of the pangenome families to infer gene acquisitions at the terminal branches. We then quantified how many times an acquisition of the same gene family of the pangenome (*i*.*e*. co-gains) occurred independently in genomes of the same CLT. This number was compared to the expected number given by a null model where the CLT does not impact the gene flow. The distribution of the expectation of the null model was made by simulation in R, taking into account the phylogeny and the distribution of CLTs. In each simulation, we used the species tree to randomly redistribute the CLT trait on the terminal branches (keeping the frequencies of CLTs equal to those of the original data). We ran 1000 simulations and compared them with the observed values with a one-sample Z-test (*75*):

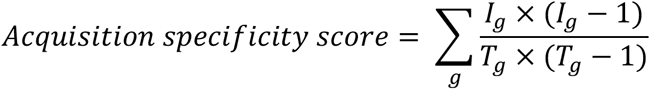

where the numerator is the number of pairs with gains in a CLT and the denominator is the number of all possible pairs. With each gene family of the pangenome *g*, the number of gene gains in strains of the same CLT *I*, and the total number of gene gains *T*. This corresponds to the sum of total number of co-gains within a CLT, normalized by the total number of co-gains for each gene. This score captures the amount of gene acquisition that happened within strains of the same CLT. If the observed score is significantly different than the simulations assuming random distribution, it means there was more genetic exchange within CLT groups than expected by chance.

### CLT specificity

We used the ancestral reconstruction of the acquisition of prophages and conjugative systems to count the number of distinct CLTs in which such an acquisition occurred. For example, one prophage family can be composed of 10 members, coming from five distinct infection events in the tree: two in KL1 bacteria and three in KL2 bacteria. Therefore, we count five acquisitions in two CLTs. The null model is that of no CLT specificity. The distribution of the expected number of CLT infected following the null model was generated by simulation (n=1000), as described above (see Identification of co-gains), and we plotted the specificity score following:

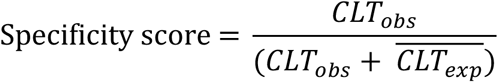

where *CLT*_*obs*_ is the observed number of serotype infected and 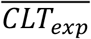 is the mean number of serotype infected in the simulations. Thus, the expected value under non-specificity is 0.5.

### Handling of draft assemblies

Since more than 90% of our genome dataset is composed of draft assemblies, *i*.*e*. genomes composed of several chromosomal contigs, we detail here the steps undertaken to reduce the impact of such fragmentation on our analysis. We only included prophages and conjugative systems that are localized on the same contig (See Prophage detection, and Analysis of conjugative systems). Kaptive is able to handle draft assemblies, and adjust the confidence score accordingly when the capsule locus is fragmented, so we relied on the Kaptive confidence score to annotate the CLT, which was treated as missing data in all the analysis when the score was below “Good” (See Capsule locus typing). For the detection of missing capsular genes, performed by Kaptive, we verified that only non-fragmented capsular loci are included (See Identification of capsule pseudogenes and inactive capsule loci). For the detection of capsule pseudogenes, we included all assemblies, and flagged pseudogenes that were localized on the border of a contig (last gene on the contig). Out of the 502 inactivated/missing genes, 47 were localized at the border of a contig. We repeated the analysis presented on figure 3B after removing these pseudogenes, and found an even better fit for the linear model at R^2^=0.77 and p=0.004. Of note, such contig breaks are likely due to IS insertions, forming repeated regions that are hard to assemble, so we kept them in the main analysis.

### Analyses of lab-evolved non-capsulated clones

To pinpoint the mechanisms by which a diverse set of strains became non-capsulated, we took advantage of an experiment performed in our lab and described previously in (*36*). Briefly, three independent overnight cultures of eight strains (Table S2) were diluted 1:100 into 5mL of fresh LB and incubated at 37°C under agitation. Each independent population was diluted 1:100 into fresh LB every 24h for three days (approximately 20 generations). We then plated serial dilutions of each population. A single non-capsulated clone per replicate population was isolated based on their translucent colony morphology, except in two replicate populations where all colonies plated were capsulated. We performed DNA extraction with the guanidium thiocyanate method (*76*), with modifications. DNA was extracted from pelleted cells grown overnight in LB supplemented with 0.7mM EDTA. Additionally, RNAse A treatment (37°C, 30min) was performed before DNA precipitation. Each clone (n=22) was sequenced by Illumina with 150pb paired-end reads, yielding approximately 1 Gb of data per clone. The reads were assembled with Unicycler v0.4.4 (*77*) and the assemblies were checked for pseudogenes (See Identification of inactive capsular loci).

### Generation of capsule mutants

Isogenic capsule mutants were constructed by an in-frame deletion of *wcaJ* by allelic exchange as reported previously (*36*). Deletion mutants were first verified by Sanger, and Illumina sequencing revealed that there were no off-target mutations.

### Conjugation assay

#### (i) Construction of pGEM-Mob plasmid

A mobilizable plasmid named pGEM-Mob was built by assembling the region containing the origin of transfer of the pKNG101 plasmid (*78*) and the region containing the origin of replication, kanamycin resistance cassette, and IPTG-inducible *cfp* from the pZE12:CFP plasmid (*79*) (see Table S3, and plasmid map, Figure S6). We amplified both fragments of interest by PCR with the NEB Q5 high-fidelity DNA polymerase, with primers adapted for Gibson assembly designed with Snapgene, and used the NEB HiFi Builder mix following vendor’s instructions to assemble the two fragments. The assembly product was electroporated into electro-competent *E. coli* DH5α strain. KmR colonies were isolated and correct assembly was checked by PCR. Cloned pGEM-Mob plasmid was extracted using the QIAprep Spin Miniprep Kit, and electroporation into the donor strain *E. coli* MFD λ-pir strain (*80*). The primers used to generate pGEM-Mob are listed in Table S4.

#### (ii) Conjugation assay

Recipient strains of *Klebsiella spp*. were diluted at 1:100 from a Luria-Bertani (LB) overnight into fresh LB in a final volume of 3mL. Donor strain *E. coli* MFD λ-pir strain (diaminopimelic acid (DAP) auxotroph) which is carrying the pGEM-Mob plasmid, exhibited slower growth than *Klebsiella* strains and was diluted at 1:50 from an overnight into fresh LB + DAP (0.3mM) + Kanamycin (50μg/ml). Cells were allowed to grow at 37° until late-exponential growth phase (OD of 0.9-1) and adjusted to an OD of 0.9. The cultures were then washed twice in LB and mixed at a 1:1 donor-recipient ratio. Donor-recipient mixes were then centrifuged for 5 min at 13,000 rpm, resuspended in 25μL LB+DAP, and deposited on a MF-Millipore™ Membrane Filter (0.45 µm pore size) on non-selective LB+DAP plates. The mixes were allowed to dry for 5 min with the lid open, and then incubated at 37°. After 1 hour, membranes were resuspended in 1mL phosphate buffered saline (PBS) and thoroughly vortexed. Serial dilutions were plated on selective (LB+Km) and non-selective (LB+DAP) plates to quantify the number of transconjugants (T) and the total number of cells (N). The conjugation efficiency was computed with:

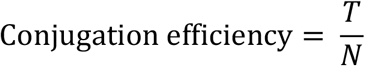

This simple method is relevant in our experimental setup because the plasmid can only be transferred from the donor strain to the recipient strain, and the duration of the experiment only allowed for minimal growth. The lack of the conjugative machinery of RK2 in the plasmid and in the recipient strains prevents transfer across recipient strains.

### Data analysis

All the data analyses were performed with R version 3.6 and Rstudio version 1.2. We used the packages ape v5.3 (*82*), phangorn v2.5.5 (*83*), and treeio v1.10 (*84*) for the phylogenetic analyses. Statistical tests were performed with the base package stats. For data frame manipulations and simulations, we also used dplyr v0.8.3 along with the tidyverse packages (*85*) and data.table v1.12.8.

## ACKNOWLEDGEMENTS

We thank Rafal Mostowy, Jorge Moura de Sousa, Nienke Buddlelmeijer, Olivier Tenaillon, Marie Touchon, and other lab members for fruitful discussions. We thank Christiane Forestier and Damien Balestrino for providing the pKNG101 plasmid and Jean-Marc Ghigo and Christophe Beloin for the gift of pZE12::CFP used to construct pGEM-Mob and *E. coli* MFD λ-pir.

## Supplementary Datasets

### Dataset SD1. Data used in this study

Genomes included in the study and their annotations. The capsule locus type (K_serotype), LPS type (O_serotype), Sequence Type (ST), N50, L90, genome size and number of contigs are indicated for each assembly, along with the total length of plasmid DNA (mlplasmid _plasmid_size), number of conjugative elements (n_CONJ) detailed in each MPF type (type F, G, I, T) and number of contigs matching at least one sequence of the PlasmidFinder database (n_plasmidfinder). We also included the inferred gene gains (gene_gains) and losses (gene_losses) for each terminal branch corresponding to the assemblies. Finally, the number of capsular pseudogenes and missing genes (n_capsular_pseudogene) and the state of the capsule locus inferred are included.

### Dataset SD2. Capsular pseudogenes table

List of capsular pseudogenes identified in the assemblies. We detail for each pseudogene the type (snp, insertion, deletion) and subtype of each mutation (frameshift, stop, complete gene deletion).

## Supplementary Tables

**Table S1.**
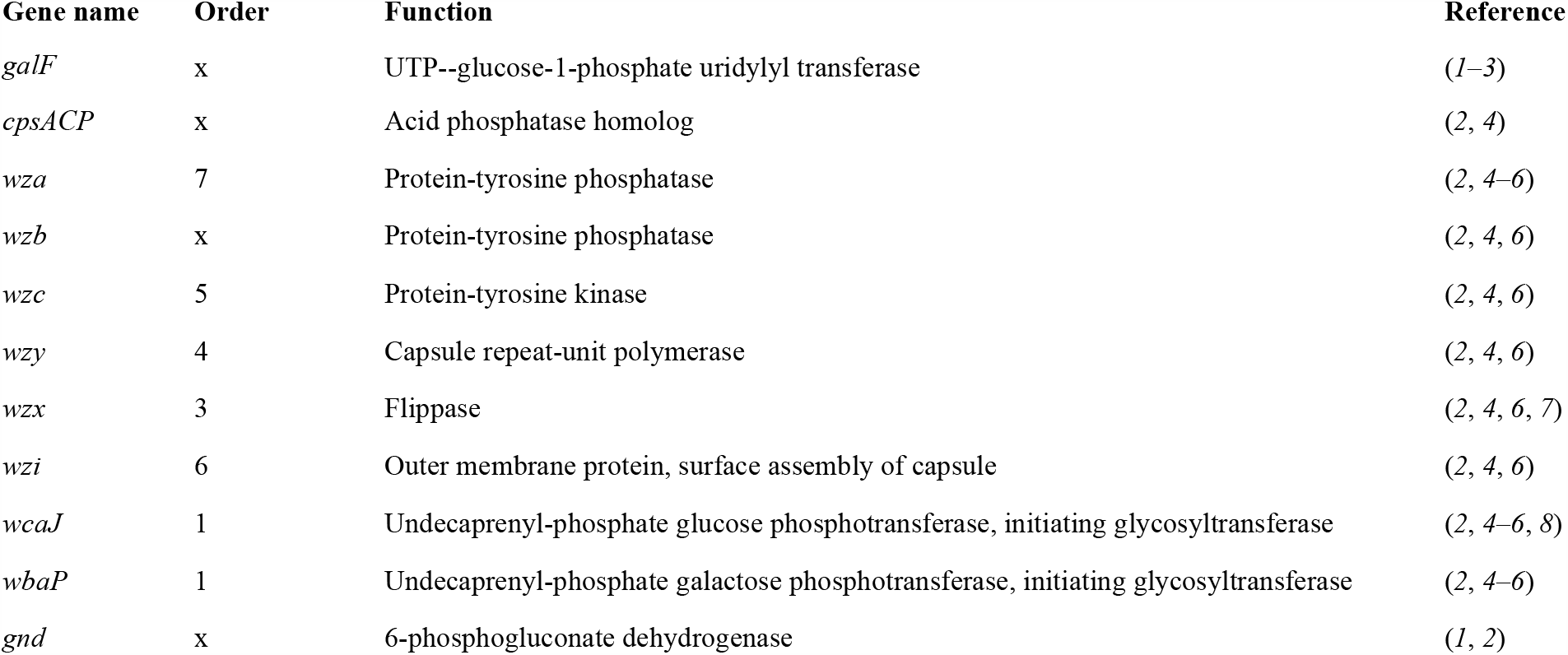
Essential genes for capsule production. The order corresponds to the order in the biosynthesis chain. The references correspond to experimental evidence that these genes are essential for capsule production.

| Gene name | Order | Function | Reference |
| --- | --- | --- | --- |
| *galF* | x | UTP--glucose-1-phosphate uridylyl transferase | (1–3) |
| *cpsACP* | x | Acid phosphatase homolog | (2, 4) |
| *wza* | 7 | Protein-tyrosine phosphatase | (2, 4–6) |
| *wzb* | x | Protein-tyrosine phosphatase | (2, 4, 6) |
| *wzc* | 5 | Protein-tyrosine kinase | (2, 4, 6) |
| *wzy* | 4 | Capsule repeat-unit polymerase | (2, 4, 6) |
| *wzx* | 3 | Flippase | (2, 4, 6, 7) |
| *wzi* | 6 | Outer membrane protein, surface assembly of capsule | (2, 4, 6) |
| *wcaJ* | 1 | Undecaprenyl-phosphate glucose phosphotransferase, initiating glycosyltransferase | (2, 4–6, 8) |
| *wbaP* | 1 | Undecaprenyl-phosphate galactose phosphotransferase, initiating glycosyltransferase | (2, 4–6) |
| *gnd* | x | 6-phosphogluconate dehydrogenase | (1, 2) |

**Table S2.** Strains used in this study.

| Strain number | Strain name | Species | ST | Capsule serotype | O-locus serotype | Country | Isolation | Accession |
| --- | --- | --- | --- | --- | --- | --- | --- | --- |
| 24 | 342 | *K. variicola* | ST146 | KL30 | O3/O3a | USA | Corn | GCF_001913175.1 |
| 26 | BJ1 | *K. pneumoniae* | ST380 | KL2 | O1v1 | France | Liver abscess | GCF_900978065.1 |
| 56 | NTHU K2044 | *K. pneumoniae* | ST23 | KL1 | O1v2 | Taiwan | Liver abscess | GCF_000009885.1 |
| 58 | SB4454 | *K. pneumoniae* | ST86 | KL2 | O1v1 | Taiwan | Liver abscess | NC_022566,<br>NC_005249,<br>SAMEA2633716 |
| 208 | SB32 | *K. pneumoniae* | ST20 | KL111 | O3b | Germany | Blood |  |
| 210 | CIP 52.229 | *K. pneumoniae* | ST59 | KL24 | O1v1 | France | Reference strain |  |
| 212 | SB5199 | *K. pneumoniae* | ST2435 | KL30 | O1v1 | France |  |  |
| 213 | SB5701 | *K. pneumoniae* | ST2435 | KL30 | O1v1 |  |  | SAMEA5753389 |
| 100 | *E. coli* S17 MFD λpir | *E. coli* |  |  |  |  | laboratory strain |  |
| 287 | *E. coli* DH5α λpir | *E. coli* |  |  |  |  | laboratory strain |  |

**Table S3.** Plasmids used in this study.

| Plasmid name | Resistance | Reference |
| --- | --- | --- |
| pKNG101 | Tet | (9) |
| pZE12-CFP | Km | (10) |
| pGEM-Mob | Km | This study |

**Table S4.**
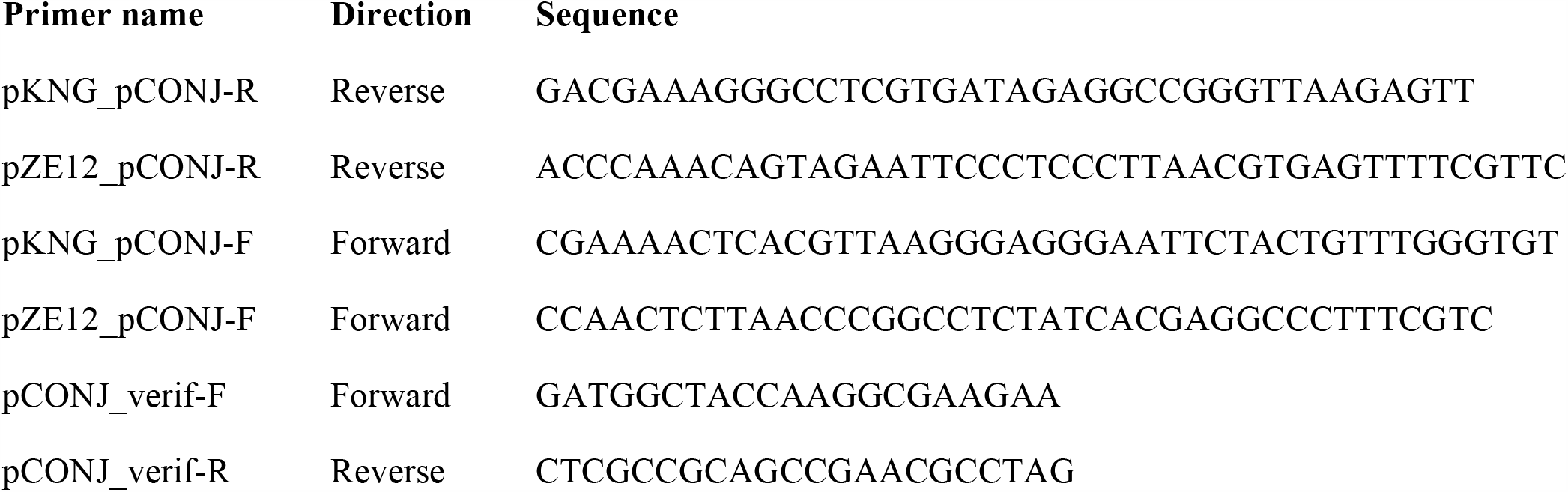
Primers used for pGEM-Mob construction.

## Supplementary figures

**Figure S1.**
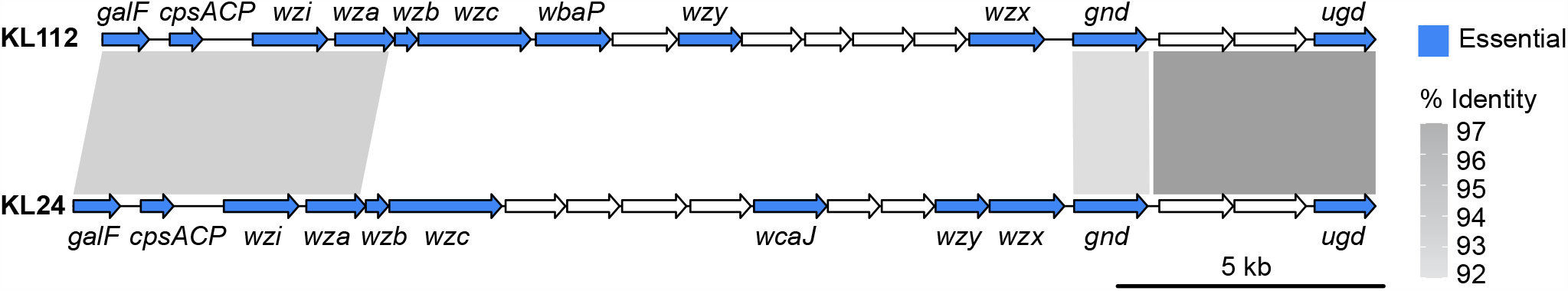
Comparison of two capsular loci types (KL112 and KL24) involved in CLT swap, with the essential genes for capsule expression colored in blue. Grey tracks correspond to the sequence identity (computed using blastn) above 90% (see scale) to indicate highly similar homologs (liable to recombine).

**Figure S2.**
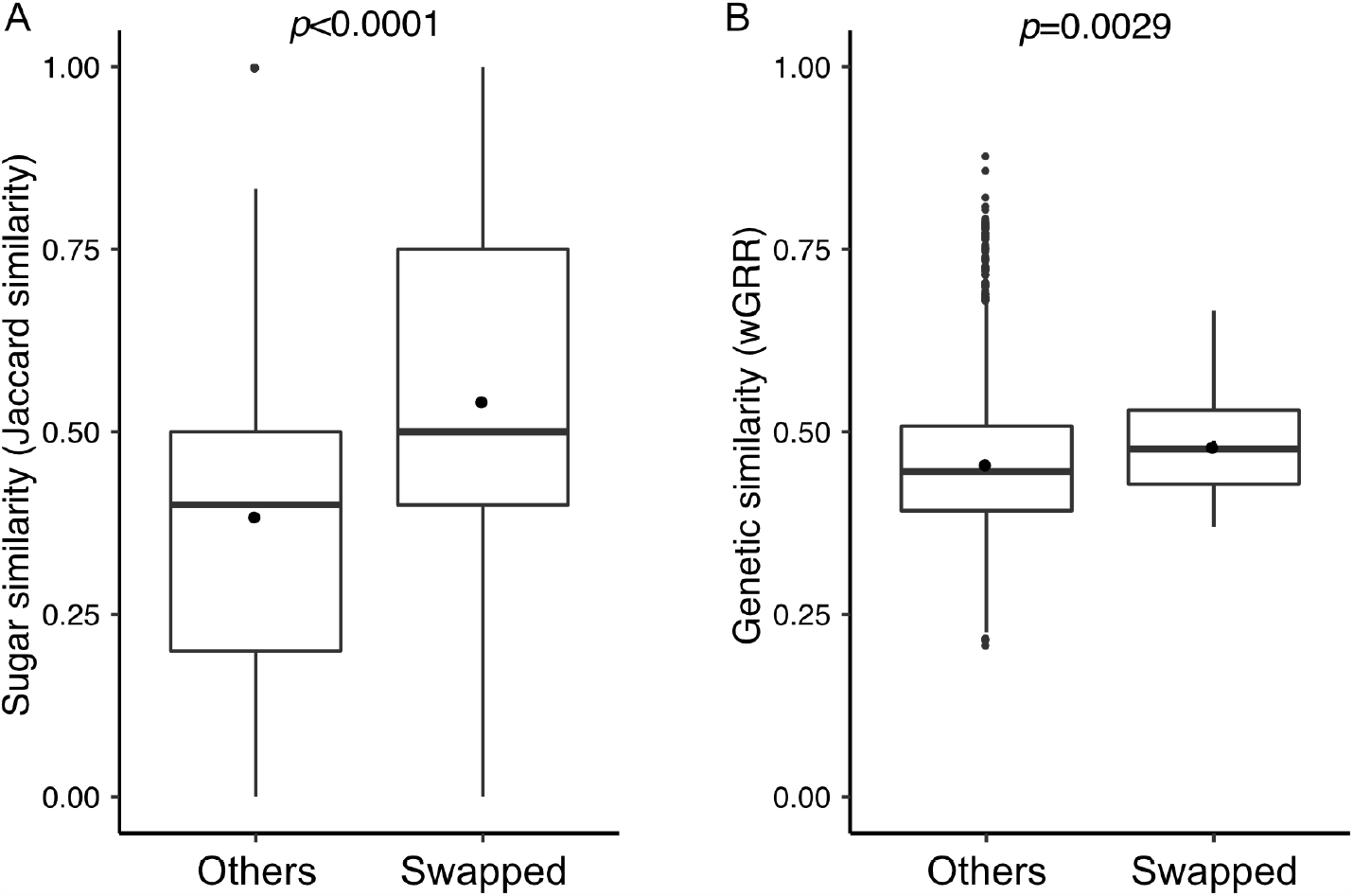
Similarity between swapped CLT and other CLT. **A**. Comparison of sugar composition similarity (Jaccard similarity) between swapped vs. others CLTs. **B**. Comparison of genetic similarity (wGRR) between swapped vs. others CLTs. The p-value displayed is for the two-sample Wilcoxon test.

**Figure S3.**
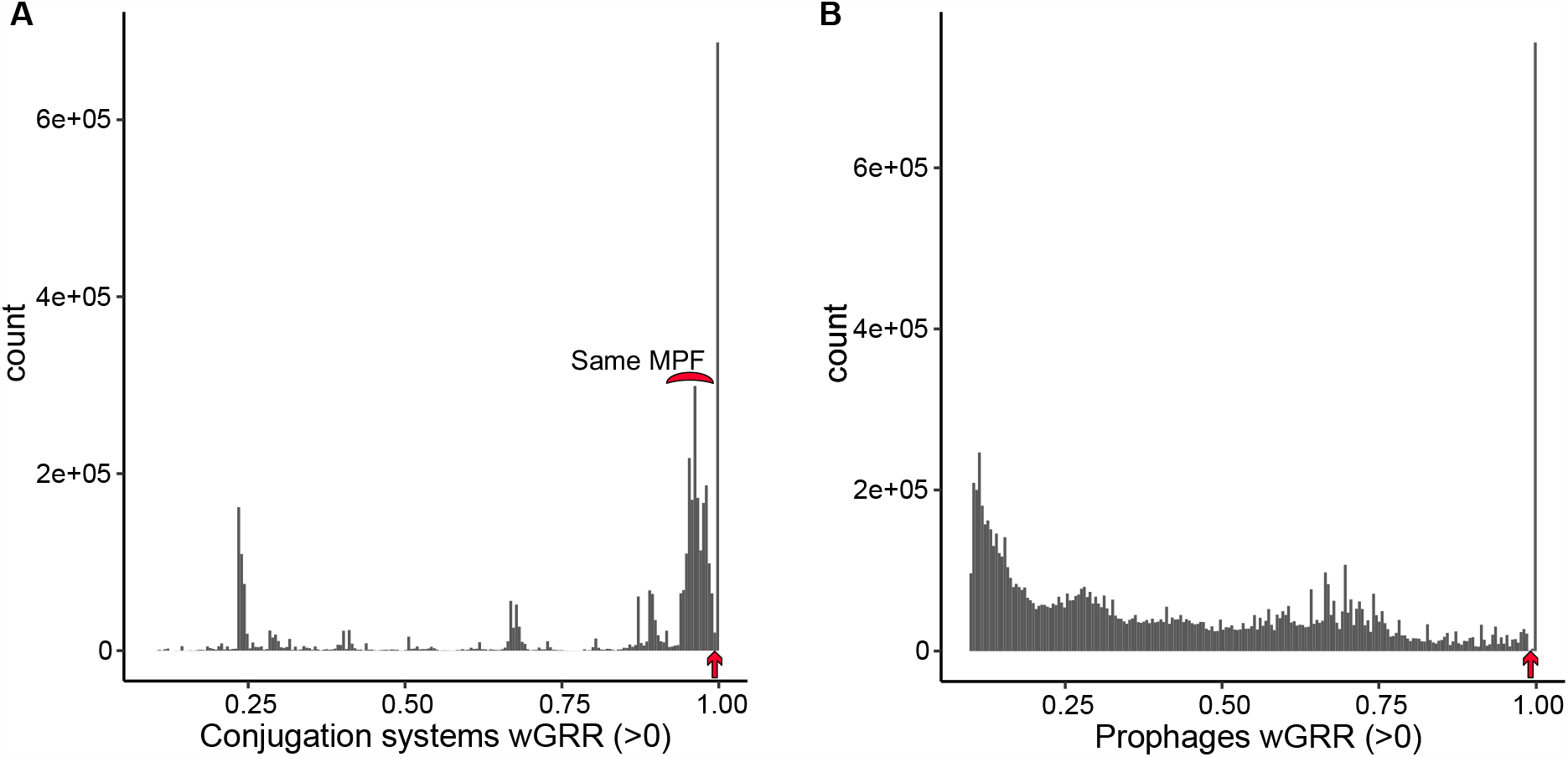
*–* Distribution of the similarity measured by wGRR between pairs of prophages (A) and between pairs of conjugative systems (B) for wGRR >0. The arrows represent the threshold (wGRR>0.99) set for clustering into families of highly similar elements. Since we performed transitive clustering to build the families, some elements belonging to the same families have wGRR<0.99. We annotated the distribution of conjugation systems belonging to the same mating-pair formation (MPF) type, which shows that systems of the same MPF are very similar but are below the selected threshold for clustering.

**Figure S4.**
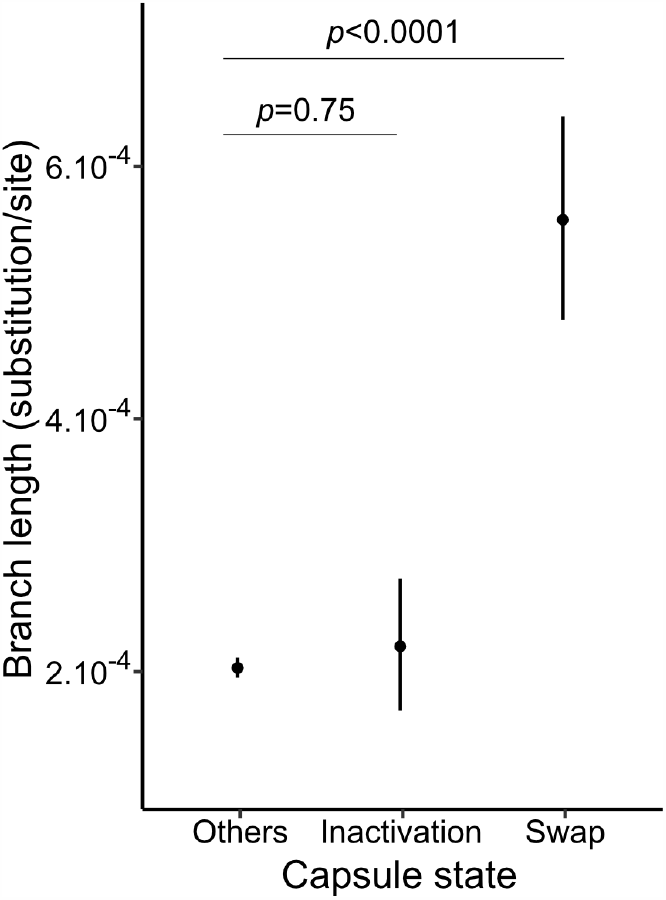
Changes in capsule state and branch length. The capsule state changes among branches of the species tree are represented on the x axis, and the branch length is represented on the y axis in substitution per site. Individual points represent the mean for each group, and the bars represent the standard error. The p-values for the t-test are represented on top of each comparisons. We also performed a two-sample Wilcoxon test to compare the medians. (“Others” vs. “inactivation”: p<0.0001. “Others” vs. “Swap”: p<0.0001)

**Figure S5.**
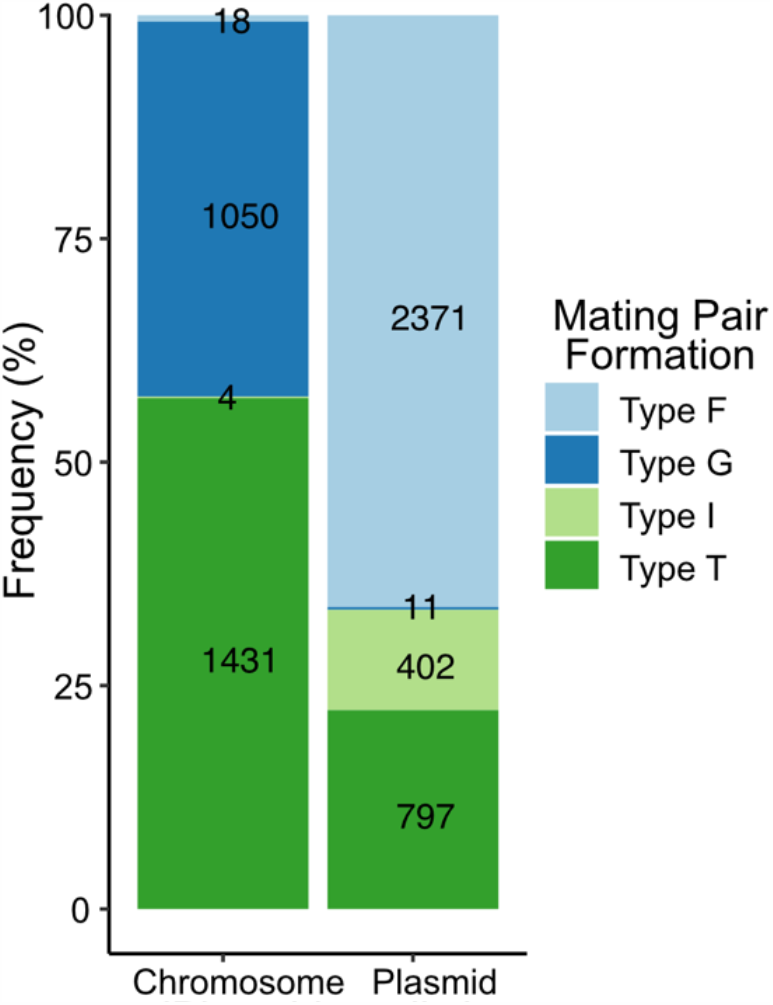
*–* Distribution of conjugation system MPF types. Conjugation systems are classified in two categories according to their genomic location, which was predicted with the mlplasmids classifier. The MPF was predicted with the CONJscan (11) module of MacSyfinder (12). Absolute number of systems are displayed for each category.

**Figure S6.**
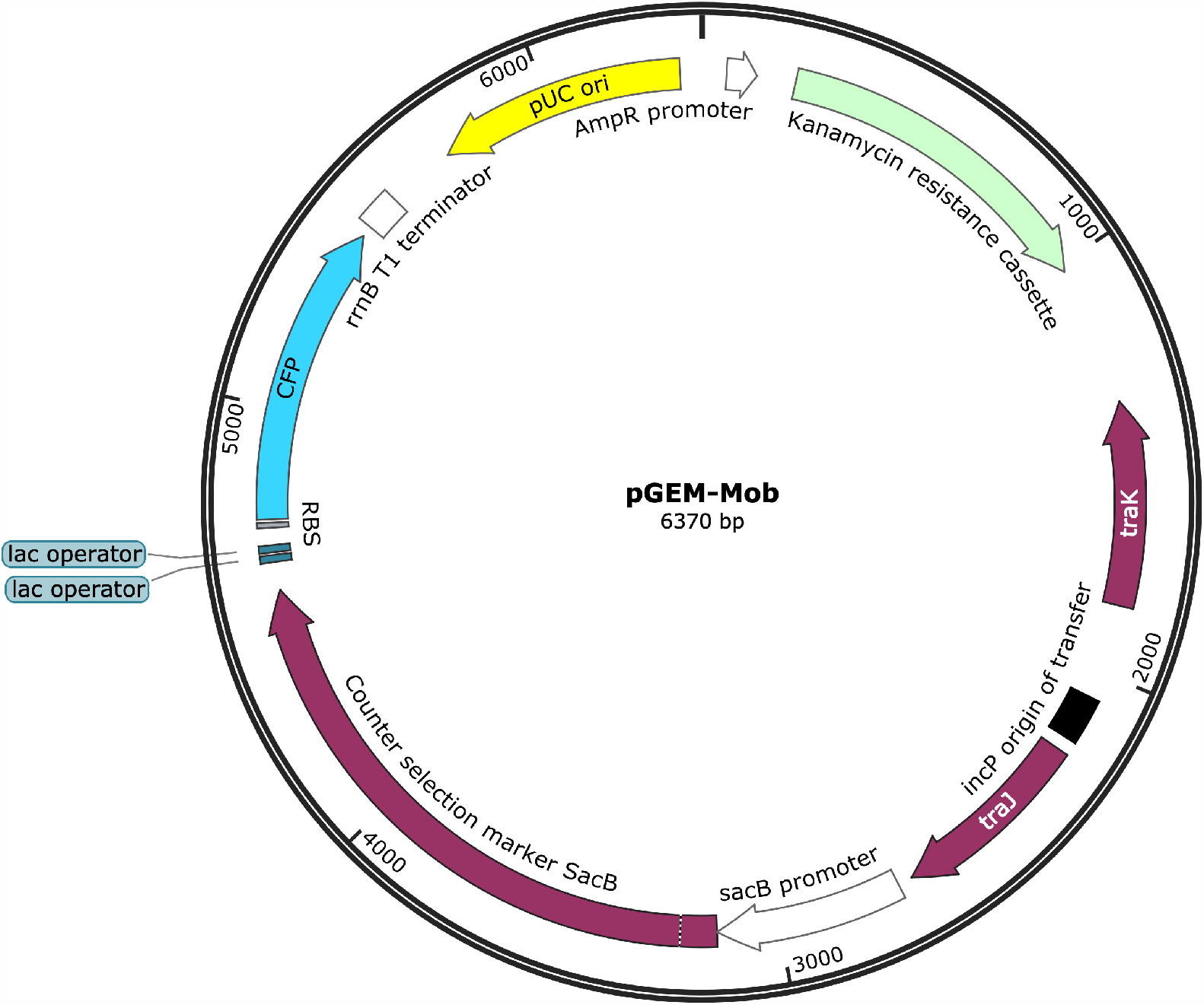
pGEM-Mob plasmid genetic map. pGEM-Mob was constructed by Gibson assembly from plasmids pKNG101 and pZE12. It encodes a colE1/pUC origin of replication (high copy number), a selectable marker (Kanamycin resistance cassette, green), the mobilizable region of pKNG101 which is composed of the origin of transfer of RK2 and two genes involved in conjugation (traJ and traK), a counter selectable marker (sacB), and an inducible CFP gene (IPTG induction). pGEM-Mob can only be mobilized in *trans* and thus can only be transferred from a strain expressing the RK2 conjugative machinery, which is absent from the panel of strains we used as recipients.

